# A *Xenopus* neuromast bioassay for chemical ototoxicity

**DOI:** 10.1101/2021.12.30.474606

**Authors:** V. Bleu Knight, Amanda R. Luna, Elba E. Serrano

## Abstract

**Background:** Ototoxic chemicals can impair the senses of hearing and balance in mammals through irreversible damage to the mechanosensory bundles of inner ear hair cells. Fish and amphibians are useful models for investigating ototoxicity because their inner ear hair cells, like those of mammals, are susceptible to damage by ototoxins. Moreover, amphibian mechanosensation is augmented by a lateral line organ on the body surface that comprises external mechanosensory hair cells. The lateral line hair cells are arranged in clusters (neuromasts) and are structurally and functionally similar to inner ear hair cells, but are more accessible for experimental manipulation. Herein, we implemented neuromasts of the amphibian (*Xenopus)* lateral line as an organ system for evaluating the effects of ototoxic chemicals, such as antibiotics, on mechanosensory hair cell bundles.

**Methods:** We examined the ultrastructure of larval *Xenopus laevis* neuromasts with scanning electron microscopy (SEM) after larvae were continuously exposed to ototoxic aminoglycoside antibiotics at sub-lethal concentrations (gentamicin; streptomycin; neomycin) for 72 hours.

**Results:** SEM images demonstrated that 72 hours of exposure to antibiotic concentrations greater than 25 µM reduced the hair cell bundle number in lateral line neuromasts.

**Conclusion:** Therapeutic drug studies will benefit from the incorporation of bioassay strategies that evaluate ototoxicity across multiple species including genera of amphibian origin such as *Xenopus*. Our outcomes support the use of the *Xenopus* lateral line for identification of potential ototoxic chemicals and suggest that *Xenopus* neuromast hair cell bundles can withstand antibiotic exposure. The *Xenopus* bioassay presented here can be incorporated into drug discovery methodology as a high-resolution phenotypic screen for ototoxic effects.

**Summary statement:** Damage to sensory cells of the inner ear by chemical agents such as antibiotics contributes to the growing global prevalence of disorders of hearing and balance. Our results demonstrate that the *Xenopus* lateral line, in conjunction with SEM, affords an accessible organ system for otoxicity screens during the drug discovery pipeline.

## 1. INTRODUCTION

Animals rely on specialized sensory cells to transmit information about mechanical stimuli to the brain. Mechanoreception by inner ear hair cells is essential for vertebrate auditory and vestibular function and is augmented in amphibians and fish through a second system, the lateral line (Fritzsch and Straka, 2014; Ghysen and Dambly, 2007). Mechanosensory inner ear hair cells share many physiological, morphological, and developmental features with hair cells of the lateral line. Regardless of species or anatomical location, hair cells are characterized by sensory bundles comprised of cilia and villi, a flask-shaped body, and innervation by the central nervous system (Fritzsch and Straka, 2014; McPherson, 2018). Lateral line hair cells form clusters, termed neuromasts, that are arranged in rows on the epidermis of fish and amphibians. For example, each neuromast of the aquatic anuran, *Xenopus laevis*, comprises between 6 and 64 hair cells that can detect particle motion in the surrounding fluid environment (Gerlach et al., 2014; Quinzio and Fabrizi, 2014; Winklbauer and Hausen, 1983). The central nervous system integrates signals from inner ear and lateral line hair cells to detect predators and prey, and to provide the organism with a sense of orientation during behavioral responses (Buck, et al., 2012; Hellman and Fritzsch, 1996; Horowitz et al., 2007; Simmons et al. 2004).

Pharmacological research has shown that ototoxins can damage or destroy mechanosensory inner ear hair cells, resulting in diminished senses of hearing and balance (Smith et al., 2016; Uribe et al., 2013; Watt, 2019). Varying degrees of ototoxic side effects have been attributed to common pharmacological agents such as antibiotics, non-steroidal anti-inflammatory drugs (NSAIDs) and antineoplastic chemicals (Yorgason et al., 2011). Among antibiotics, aminoglycosides have a lengthy history of therapeutic use, yet are widely recognized as causal factors of auditory and vestibular dysfunction (Jiang et al., 2017; Selimoglu, 2007). Studies with inner ear hair cells have provided mechanistic evidence for aminoglycoside ototoxicity, e.g. gentamicin has been shown to block hair cell mechanotransduction, and to trigger signal cascades that lead to the destruction of hair cell stereocilia and cell death through apoptosis (Coffin et al., 2013; Ernst et al., 1994; Kroese and van den Bercken, 1982; Williams et al., 1987). These investigations have sparked interest in evaluating the potential ototoxic effects of novel or established therapeutic agents. However, the sequestered location and small number of inner ear hair cells can impede the rapid analysis of drug action and has driven the search for accessible cellular alternatives (Brack and Ramcharitar, 2012; Hessel et al., 2018; Steyger et al, 2018; Vlasits et al., 2012).

The superficially located mechanosensory cells of the lateral line are an ideal complementary cellular alternative to inner ear hair cells in a drug screening pipeline. Anatomical and physiological studies have demonstrated the preservation of features of the lateral line and inner ear structures across multiple species (Pujol-Martí and López-Schier 2013; Fritz and Straka, 2014). The advantages posed by accessible lateral line neuromasts have led to their use in evaluating ototoxic and otoprotective drugs in organisms such as fish, amphibians, and salamanders (Holmgren and Sheets, 2021). The lateral line of the fish, *Dario renio* has emerged as a popular experimental system for ototoxicity investigations due to the many resources available for Zebrafish cellular and molecular analysis as well as for rapid follow-up genetic screens (Coffin et al., 2010; Domarecka et al, 2020; Holmgren and Sheets, 2021; Ou et al., 2012; Rocha-Sanchez et al., 2018; Stengel, et al., 2017; Ton and Parng, 2005).

In comparison to fish, the amphibian lateral line has been underutilized as a target for otoxicity research. Outcomes from amphibian lateral line studies enhance a body of influential research that has used amphibian inner ear organs to probe hair cell development and regeneration and the physiological process of mechanotransduction (Burns and Corwin, 2013; Hudspeth, 1989; McPherson, 2018). Among amphibians, *Xenopus* holds promise for otoxocity studies due to its well characterized sensory hair cell systems for lateral line and inner ear mechanoreception. A prominent developmental model organism, *Xenopus* has contributed to our knowledge of inner ear hair cell structure and function (Lopez-Anaya et al., 1997; Mason et al., 2009; Niewkoop and Faber, 1967; Quick and Serrano, 2005; Shin et al., 2005, Smotherman and Narins, 2000; Straka and Chagnaud, 2017; Vedurmudi et al, 2018). *Xenopus* inner ear hair cells, like those of other amphibians, are sequestered in eight endorgans that are more compartmentalized and respond to a wider and higher frequency range than those of fish (Bever et al, 2003; Paterson, 1948; Quick and Serrano, 2005).

Like their zebrafish counterparts, mechanosensory hair cells of the amphibian lateral line are superficially located and are a tractable cellular model for screening drug ototoxicity (Lum et al., 1982; Simmons et al., 2015). Moreover, structural and physiological changes to *Xenopus* lateral line hair cells following aminoglycoside exposure have been documented using electrophysiological approaches and fluorescence light microscopy (Kroese and van den Bercken, 1982; Nishikawa and Sasaki, 1996). For these reasons, the *Xenopus* lateral line is a promising system for identifying ototoxic compounds, and to verify and complement ototoxicity studies that have utilized the zebrafish lateral line.

Here we present results of our efforts to expand the utility of the *Xenopus* lateral line for ototoxin evaluation. To this end, we used scanning electron microscopy to evaluate the impact of aminoglycoside antibiotics on neuromast ultrastructure. We aimed to develop a straightforward protocol to determine whether chemical agents, such as aminoglycosides, cause specific damage to neuromast hair cell stereocilia using an imaging method that affords higher resolution of cellular detail than is achieved with fluorescence microscopy. Our data demonstrate that this whole animal bioassay can be used to screen water-soluble chemicals, such as the ototoxic aminoglycoside antibiotics, for their ability to damage the ultrastructure of *Xenopus* mechanosensory lateral line cells. The impact of our findings is augmented by the emergence of substantial international resources for *Xenopus (e*.*g. Xenbase*, the *National Xenopus Resource*, and the *European Xenopus Resource Centre)* that leverage contemporary molecular and genomics methods for mechanistic studies of disease and development in *Xenopus* (Blum and Ott, 2018; Fortriede et al, 2019; Horb et al, 2019).

## 2. MATERIALS AND METHODS

### 2.1 Experimental specimens and institutional oversight

All animal care and handling procedures were approved and overseen for compliance with federal guidelines for amphibian research by the Institutional Animal Care and Use Committee (IACUC) of New Mexico State University (*PHS Approved Animal Welfare Assurance Number* D16-00564; A4022-01). Newly fertilized embryos were ordered from NASCO (Fort Atkinson, WI) and allowed to develop until stage 46/47 (staged according to Niewkoop and Faber, 1967), typically 4 days post fertilization. Upon hatching, larvae were fed *Xenopus* Express Premium Tadpole food powder (*Xenopus* Express Inc.)

### 2.2 Antibiotic solutions

Stage 46/47 *Xenopus* were exposed to a range of concentrations of gentamicin (25µM, 50 µM, 100 µM, 200 µM, 400 µM), streptomycin (100 µM, 400 µM), or neomycin (100 µM, 400 µM). *Xenopus* were maintained at temperatures ranging from 16° C to 20° C in 4L aquaria until transfer to individual dishes for antibiotic exposure. Fresh stock solutions (50 mM) of gentamicin sulfate (Sigma-G1264, Lot#097k06883v), streptomycin sulfate (Sigma-S6501, Lot# 02k07352v), or neomycin trisulfate (Sigma-N5285, Lot#080m0048v) salts were prepared in 0.1M phosphate buffered saline and adjusted to pH (7.0-7.5) with sodium bicarbonate prior to filter sterilization. Stock solutions of antibiotic were diluted in water and the resulting antibiotic solutions were plated in the chambers of covered 6-well tissue culture trays (35 × 14 mm; 7 mL per well; 1 larvae per well). Antibiotic solutions were prepared from stock immediately prior to animal introduction.

### 2.3 Exposure protocol

The treatment groups (gentamicin: 25 µM, 50 µM, 100 µM, 200 µM, 400 µM; streptomycin: 100 µM, 400 µM; neomycin 100 µM, 400 µM; n=5) were exposed to antibiotics for a total of 72 h (Table 1). The antibiotic solution was replaced with fresh antibiotic solution at 48 h. Following the 72 hr exposure period, tadpoles from each treatment group were immediately euthanized in 0.1% tricaine methanesulfonate MS-222, (Finquel® TR12040; pH = 7) in phosphate buffered saline (PBS) containing 40 mM sodium bicarbonate and processed for scanning electron microscopy). Control groups were not exposed to antibiotic. Control group larvae were placed in 7 mL of fresh water in 6-well plates with a fresh water change after 48 h. Control group larvae (n = 5) were euthanized (MS-222) 72 h after the start of the experiment in parallel with the antibiotic-treated groups and processed for SEM.

**Table 1:** Experimental paradigm for antibiotic exposure regimen. The experimental design implemented a control group and two treatment groups that were exposed to three antibiotic regimens. Larval Control Groups were not exposed to antibiotic and were euthanized at the same time points as the treatment groups. Larvae were exposed to antibiotic for 72 h and immediately euthanized.

| Experimental Group | Exposure | Concentration |
| --- | --- | --- |
| Control | No Antibiotic | 0 μM |
| Treatment Group: Euthanized after 72 h of continuous exposure | Gentamicin | 25 μM |
| | | 50 μM |
| | | 100 μM |
| | | 200 μM |
| | | 400 μM |
| | Streptomycin | 100 μM |
| | | 400 μM |
| | Neomycin | 100 μM |
| | | 400 μM |

### 2.4 Developmental Stages and Behavior

Animals were staged at the beginning of the antibiotic exposure and after euthanasia using the criteria of Nieuwkoop and Faber (1967). The reflexive startle response of the animals was assessed by tapping the center of the 6-well plate with a pencil at the beginning of the experiment (prior to the introduction of antibiotics), at the 48 h point after antibiotics were replenished, and at the 72 h experimental end point

### 2.5 Scanning Electron Microscopy

Three larvae from each treatment group were processed for scanning electron microscopy of individual neuromasts from the anterior lateral line. Following euthanasia, specimens were rinsed in distilled water, then placed in 2.5% glutaraldehyde (Electron Microscopy Sciences-16220) in 0.1 M cacodylate buffer (pH 7.2; Electron Microscopy Sciences-11654) for a minimum of 8 h at 4°C. Samples were rinsed twice in 0.1M cacodylate buffer (pH 7.2) and twice in distilled water for 5 min. each and then dehydrated in an ethanol series (25%, 50%, 75%, 100%, 100%) for 10 minutes each. Samples were chemically dried by placing them on a filter paper soaked with Hexamethyldisilazane (HMDS) solvent in a Petri-dish overnight in a fume hood. The specimens were mounted on SEM stubs with double-sided sticky tape, then loaded onto a Denton vacuum evaporator (Denton Desk IV Sputter Coater) and sputter coated with a thin layer of gold. Individual lateral line neuromasts were imaged with a Hitachi S-3400N SEM (0.3 - 30 kV).

### 2.6 Data Analysis

The number of visible hair cell bundles in individual neuromasts was quantified from SEM images of the posterior lateral line (PLL) and anterior lateral line (ALL) of the animals in each experimental condition. SEM images were manually analyzed using a “blind” protocol where the experimental condition was unknown during the tally of ciliary bundles. Bundles were counted in neuromasts when more than one visible stereocilium and/or kinocilium emerged in a group or cluster from the surface of the neuromast. The number of bundles per neuromast was averaged for 4-7 neuromasts per larva. We noted that sometimes neuromasts could not be located on the lateral line of SEM images regardless of treatment condition. This average was used to compute the average for each experimental condition (n = 5 larvae per group; 4-7 neuromasts per larva). Statistically significant differences between experimental conditions were calculated using a mixed-model nested analysis of variance ANOVA with the Satterthwaite approximation, followed by the Tukey test (McDonald, 2009).

## 3. RESULTS

### 3.1 Behavioral and developmental changes were not detected in response to antibiotic exposure

*Xenopus* larvae in experimental groups (Table 1) were staged according to Nieuwkoop and Faber (1967) prior to the start of the experiment and upon euthanasia. Treatment groups comprised *Xenopus* larvae exposed to antibiotic (gentamicin: 25 µM, 50 µM, 100 µM, 200 µM, 400 µM; streptomycin: 100 µM, 400 µM; neomycin 100 µM, 400 µM) for 72 h and immediately euthanized. *Xenopus* larvae in the control group were euthanized at the same time point. Differences in larval development were not detected between control and treatment groups during the four-day time frame of this experiment. At the end of the experiment, specimens had progressed from stages 46/47 to stages 47/48 in all experimental groups. There was no mortality among control or treatment groups during the exposure regimen.

The startle response (where the larvae swim away from the source of the disturbance) is partially mediated by lateral line hair cells and can be used as an indicator of viable mechanosensation in an organism (Buck et al., 2012; Hailey et al., 2012). A startle response was elicited in control and treatment groups prior to antibiotic exposure at the start of the experiment, at the 48 h point, and after 72 h of antibiotic exposure in order to evaluate the functionality of mechanosensation pathways. Differences were not detected between control and treatment groups when startle response behaviors were assessed at the designated time points.

### 3.2 Antibiotic exposure alters neuromast bundle ultrastructure

High-resolution images of individual lateral line neuromasts acquired with scanning electron microscopy, which permitted the evaluation of the structural integrity of neuromast hair cell bundles from larvae following a 72 h treatment with gentamicin (Fig. 1; 25, 50, 100 200, 400 µM); streptomycin (Fig 2; 100, 400 µM); or neomycin (Fig 2; 100, 400 µM). Micrographs of neuromasts from animals treated with gentamicin, streptomycin or neomycin revealed ultrastructural differences between the hair cell bundles of control and treatment group neuromasts (Figs. 1 and 2). Qualitative observations suggest that stereocilia and/or kinocilia in hair cell bundles of treatment groups, when present, appeared reduced in length. Stereocilia and/or kinocilia length appeared to be less affected by neomycin exposure than by exposure to streptomycin or gentamicin. Manual and digital quantification of bundle length were precluded by the convoluted, collapsed, and intertwined appearance of the ciliary bundles in many neuromast images (see Figs. 1 and 2 for examples).

**Figure 1:**
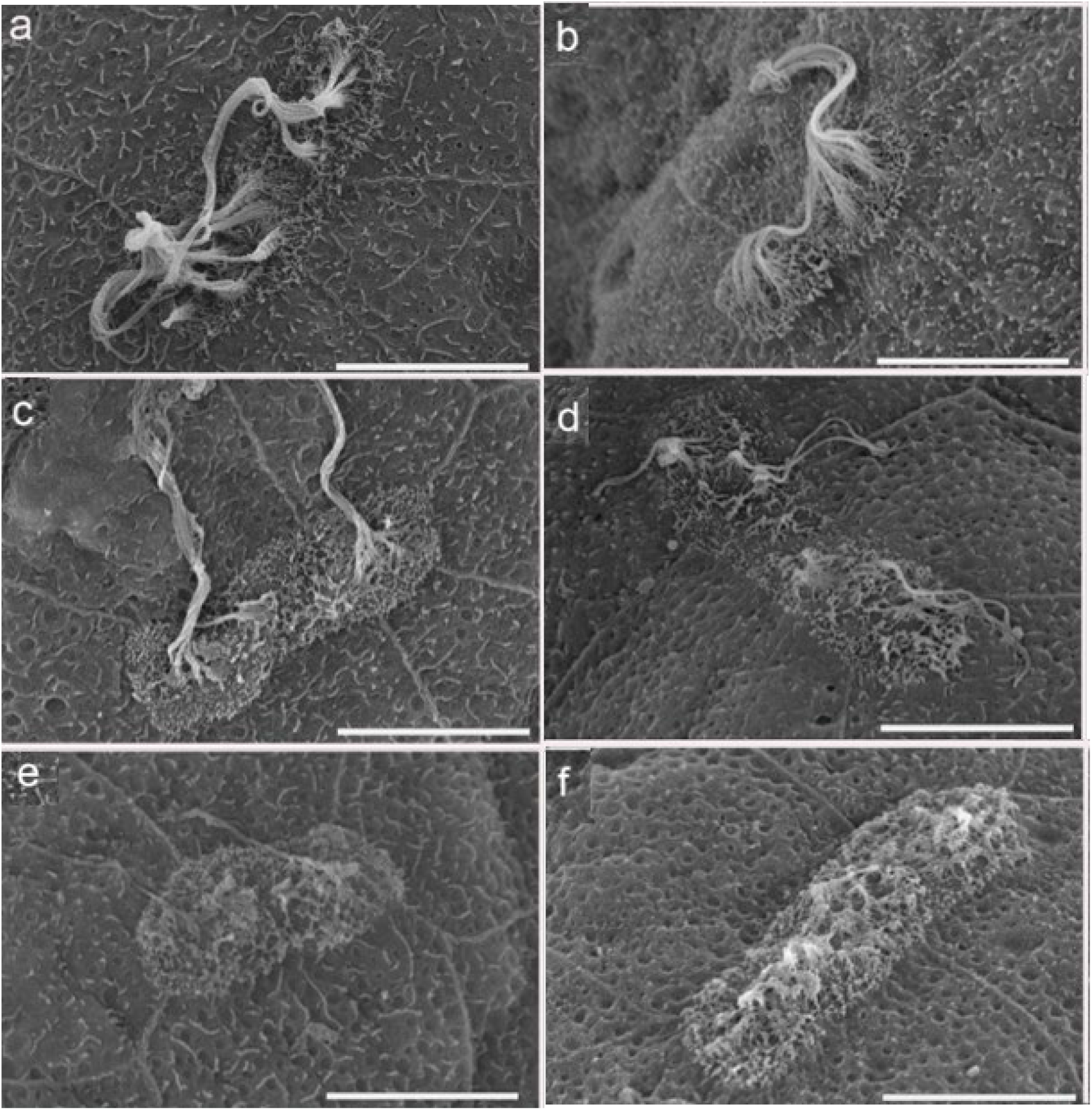
Imaging of neuromast hair cell bundles exposed to gentamicin: Representative SEM images of lateral line neuromasts from control animals (a) and larvae continuously exposed to gentamicin (25 µM, b; 50 µM, c; 100 µM, d; 200 µM, **e**; 400 µM, k) for 72 hours. Scale bar = 10µm.

**Figure 2:**
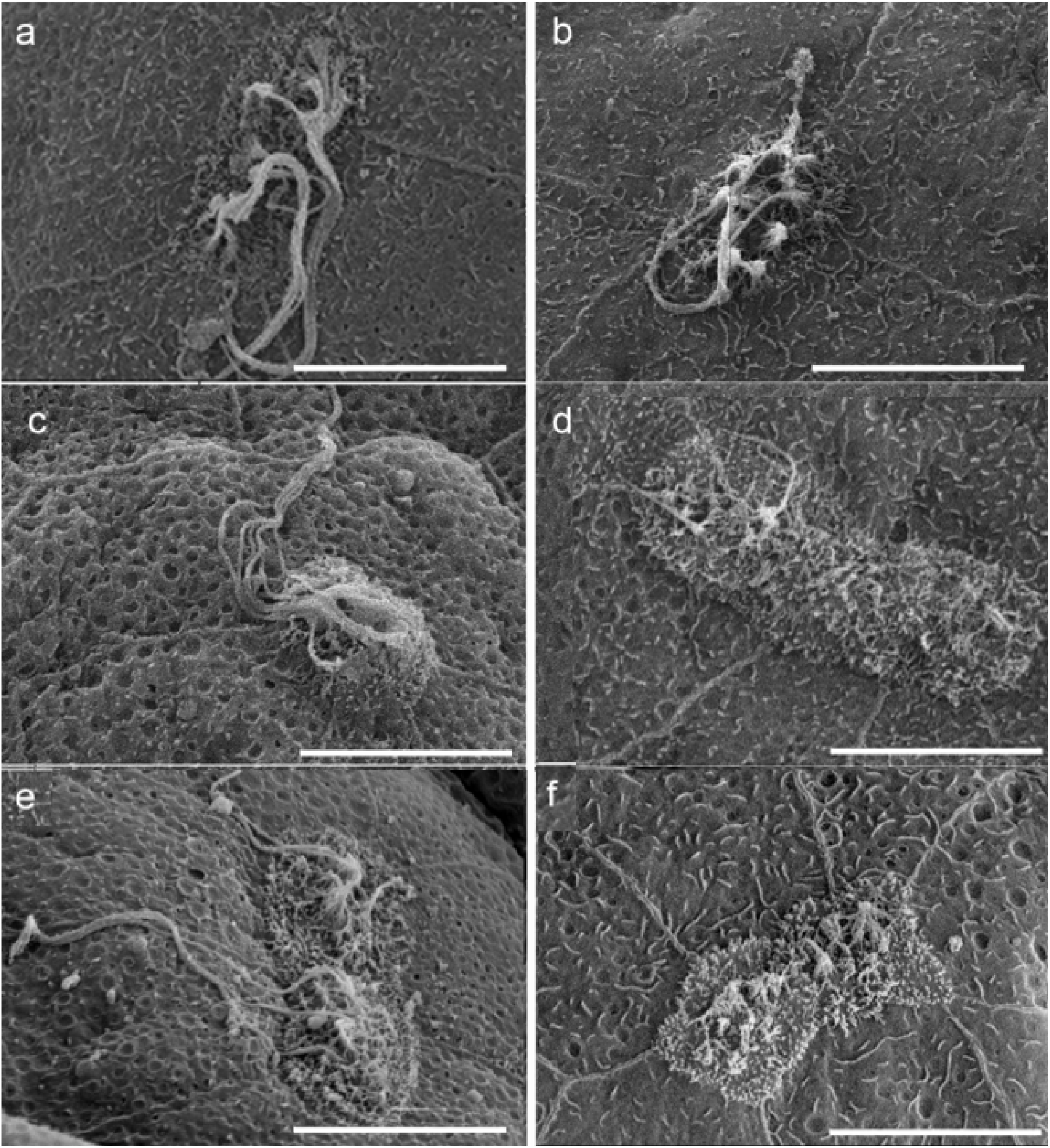
Imaging of neuromast hair cell bundles exposed to neomycin and streptomycin: Representative SEM images of lateral line neuromasts from control animals (a, b) and larvae continuously exposed to neomycin (c; 100 µM, d; 400 µM) streptomycin (e; 100 µM, f; 400 µM) for 72 hours. Scale bar = 10µm.

### 3.3 Antibiotic exposure affects hair cell bundle numbers

SEM images were analyzed to determine whether antibiotic exposure damaged neuromasts by altering the number of hair cell bundles per neuromast. Hair cell bundles comprising more than one stereocilia emerging in a group or cluster from the surface of the tadpole were tallied for individual neuromasts (Figs. 1-3). In control stage 47/48 *Xenopus* larvae, neuromasts were populated by an average of 5.4 ± 0.4 and (mean ± S.E.) hair cell bundles.

**Figure 3:**
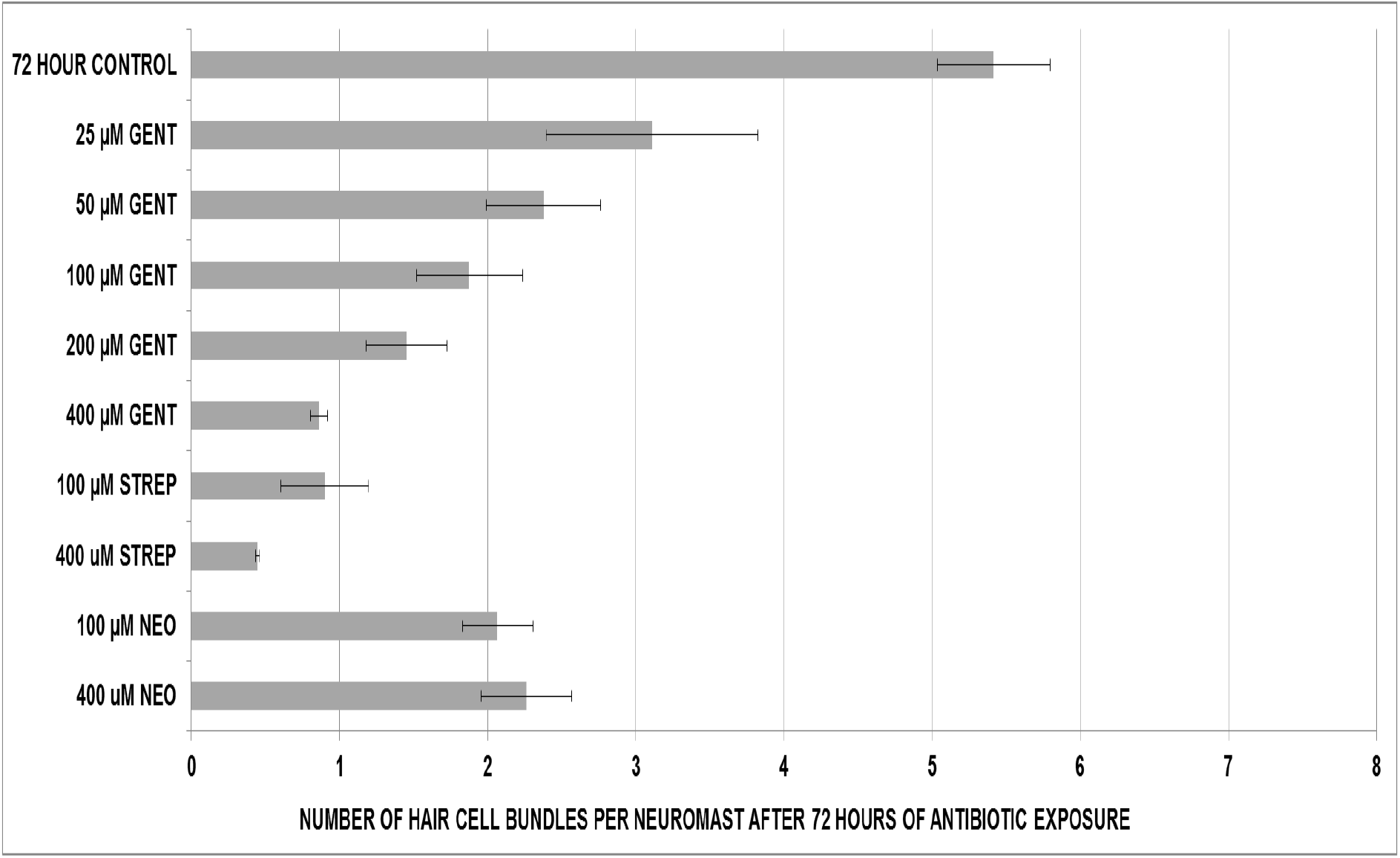
Antibiotic effects on hair cell bundles: The average number of hair cell bundles per neuromast is shown for control larvae (top bar) and after 72 hours of treatment with varying concentrations of gentamicin (GENT), streptomycin (STREP), or neomycin (NEO). Results are displayed as mean ± S.E. (n = 3).

Two statistical methods were used to assess whether antibiotic exposure altered the number of neuromast hair cell bundles immediately after an exposure period, and after antibiotic removal. A multi-level nested ANOVA analysis was undertaken to evaluate the null hypothesis, that there is no difference between the numbers of hair cell bundles per neuromast in all experimental groups. Outcomes of the ANOVA indicate a p value of 5.36 × 10^−6^.

The Tukey test was used to evaluate the extent to which experimental groups differed from one another (Table 2). We noted that mean values for each experimental group met Tukey significance threshold criteria of 99% confidence, α = 0.01 if they differed by at least 2.67 bundles per neuromast. The Tukey test revealed that after 72 hours of continuous exposure, animals treated with concentrations of gentamicin, streptomycin, and neomycin greater than or equal to 50 µM had fewer hair cell bundles per neuromast as compared to the control group at 72 hours, and also met the Tukey criteria.

**Table 2:**
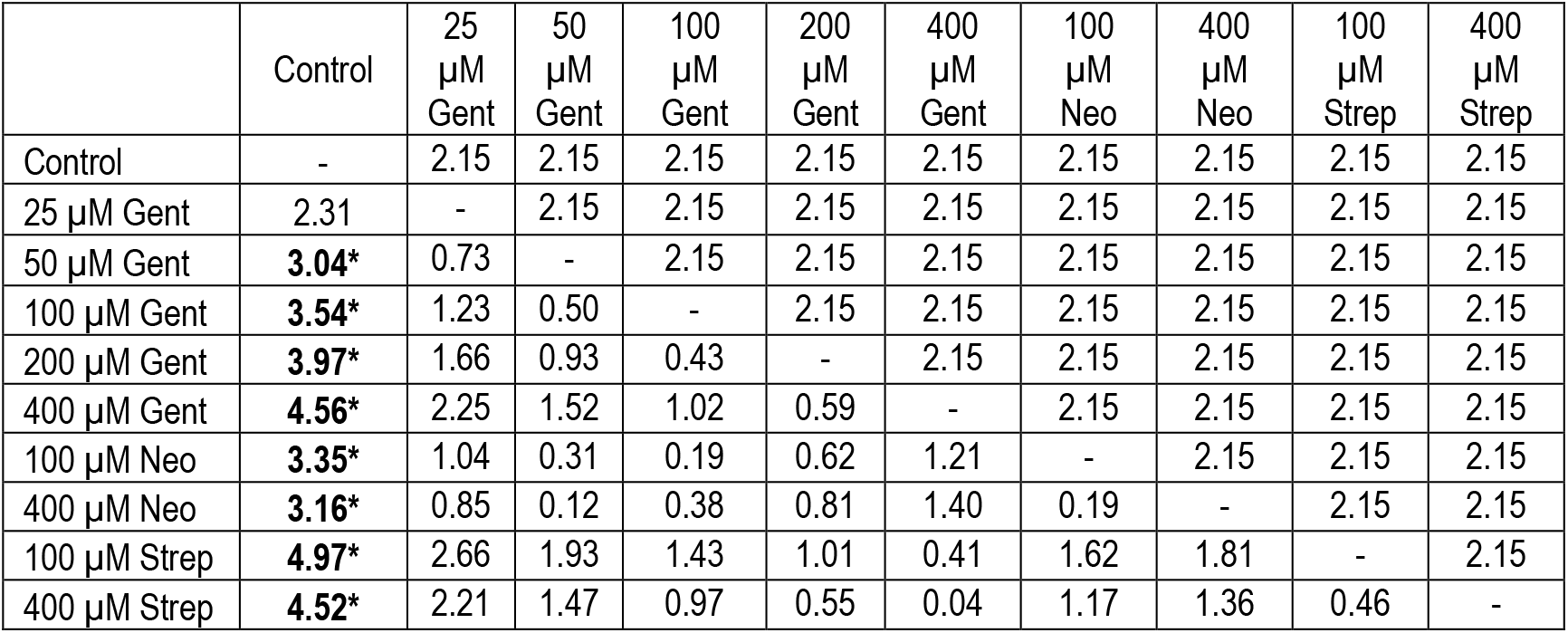
Tukey analysis of differences between group means. Table showing experimental group means and differences between means for experimental groups. Differences between means based on the Tukey-Kramer minimum significant difference of 2.67 (α = 0.01; 99% confidence) are marked with an * and displayed in bold print for comparisons.

| | Control | 25<br>$\mu\text{M}$<br>Gent | 50<br>$\mu\text{M}$<br>Gent | 100<br>$\mu\text{M}$<br>Gent | 200<br>$\mu\text{M}$<br>Gent | 400<br>$\mu\text{M}$<br>Gent | 100<br>$\mu\text{M}$<br>Neo | 400<br>$\mu\text{M}$<br>Neo | 100<br>$\mu\text{M}$<br>Strep | 400<br>$\mu\text{M}$<br>Strep |
| --- | --- | --- | --- | --- | --- | --- | --- | --- | --- | --- |
| Control | - | 2.15 | 2.15 | 2.15 | 2.15 | 2.15 | 2.15 | 2.15 | 2.15 | 2.15 |
| 25 $\mu\text{M}$ Gent | 2.31 | - | 2.15 | 2.15 | 2.15 | 2.15 | 2.15 | 2.15 | 2.15 | 2.15 |
| 50 $\mu\text{M}$ Gent | **3.04\*** | 0.73 | - | 2.15 | 2.15 | 2.15 | 2.15 | 2.15 | 2.15 | 2.15 |
| 100 $\mu\text{M}$ Gent | **3.54\*** | 1.23 | 0.50 | - | 2.15 | 2.15 | 2.15 | 2.15 | 2.15 | 2.15 |
| 200 $\mu\text{M}$ Gent | **3.97\*** | 1.66 | 0.93 | 0.43 | - | 2.15 | 2.15 | 2.15 | 2.15 | 2.15 |
| 400 $\mu\text{M}$ Gent | **4.56\*** | 2.25 | 1.52 | 1.02 | 0.59 | - | 2.15 | 2.15 | 2.15 | 2.15 |
| 100 $\mu\text{M}$ Neo | **3.35\*** | 1.04 | 0.31 | 0.19 | 0.62 | 1.21 | - | 2.15 | 2.15 | 2.15 |
| 400 $\mu\text{M}$ Neo | **3.16\*** | 0.85 | 0.12 | 0.38 | 0.81 | 1.40 | 0.19 | - | 2.15 | 2.15 |
| 100 $\mu\text{M}$ Strep | **4.97\*** | 2.66 | 1.93 | 1.43 | 1.01 | 0.41 | 1.62 | 1.81 | - | 2.15 |
| 400 $\mu\text{M}$ Strep | **4.52\*** | 2.21 | 1.47 | 0.97 | 0.55 | 0.04 | 1.17 | 1.36 | 0.46 | - |

## 4. DISCUSSION

Modern medicine has extended the human life span and enhanced the quality of life, in large part due to the discovery of therapeutic agents such as antibiotics. However, chemical interventions frequently have unforeseen and undesirable consequences. For example, the aminoglycoside antibiotics (e.g. gentamicin, neomycin, streptomycin) were among the first drugs prescribed over 60 years ago as treatments for bacterial infections. Although these antibiotics are powerful treatments against sepsis and other infections, their deleterious side effects include nephro- and ototoxicity (Boyer et al., 2013; Selimoglu, 2007; Watt, 2019). Aminoglycoside ototoxicity results in hearing loss and decreased vestibular function that is frequently attributed to damaged mechanosensory hair cells (Kroese and van den Bercken, 1982; Selimoglu, 2007).

The prevalence of hearing loss and vestibular disorders is increasing globally, yet the extent to which established or novel drugs can damage mechanosensory cells remains undetermined (Chiu et al., 2008; Kochkin, 2009; Olusanya et al, 2019). The ability to rapidly screen new compounds for potential ototoxic side effects may facilitate the development of therapeutic agents that can protect, or do not compromise, hearing and balance. The preclinical research phase of the drug development pipeline relies on screens with cell line and animal models that share molecular, cellular, and physiological properties with human cells and organs. Efforts to establish cell lines that produce fully differentiated mechanosensory hair cells with their characteristic stereocilia and mechanotransduction channels have been unsuccessful and investigations rely on the use of animal tissues and organoids (Koehler et al. 2017; Lim et al, 2018).

The *Xenopus* lateral line bioassay presented here is intended to expand upon and complement data collected from other non-human animal models as part of ototoxicity screening protocols. The lateral line is a readily accessible epidermal specialization of fish and aquatic amphibians comprising neuromast receptor organs consisting of mechanosensory hair cells surrounded by supporting cells (Piotrowski and Baker, 2014). In these species, the perception of fluid motion depends upon the integration of information from mechanosensory hair cells in lateral line neuromasts and the inner ear (Buck et al., 2012; Coffin et al., 2010; Hellmann and Fritzsch, 1996; Horowitz et al., 2007).

The anatomy and physiology of the lateral line offer advantages for high throughput investigations because chemical agents can be delivered in water, allowing investigators to evaluate hair cell function *in vivo* through the assessment of a behavioural phenotype such as the startle response and rheotactic (Simmons et al., 2004). Neuromast hair cells share developmental, morphological, and physiological features with inner ear hair cells, such as a mechanosensory bundle comprising stereocilia, synaptic vesicle release, and neural innervation (Atkinson et al., 2015; Quinzio and Fabrezi, 2015). Lateral line hair cells from *Xenopus laevis* have been used to evaluate the physiological effects of aminoglycosides (Kroese and van den Bercken, 1982; Nishikawa and Sasaki, 1996). However, the ultrastructure of hair cell stereocilia after antibiotic exposure has not been thoroughly investigated.

Our experiments were undertaken with neuromasts from *X. laevis* larval stage 46/47 due to the deceleration of developmental progression compared with earlier stages, and the relatively constant number and size of the neuromasts that are characteristic of this stage (Nieuwkoop and Faber, 1967; Winklbauer, 1989). At developmental stage 46/47, the lateral lines have differentiated into individual sense organs, and the lateral line neuromasts are not visually obstructed by the cupulae. At this stage, the inner ear’s vestibular sensory structures (canals, utricle, sacculus) are undergoing early compartmentalization, while the auditory organs are not yet formed (Quick and Serrano, 2005; Winklbauer, 1989).

The antibiotic concentrations (25-400 µM) used in our experimental paradigm are estimated to represent concentrations of antibiotics that could be found in human tissues. Therapeutic doses of intravenous gentamicin given to humans raise the concentration in blood and serum to approximately 20 µM (Glew et al., 1976; Setiabudy et al., 2013). However, concentrations of gentamicin in organs where aminoglycosides have toxic action may be significantly more than this amount. For example, gentamicin can be measured in the endolymph for up to 15 days after the cessation of treatment, whereas it disappears from plasma and perilymph much faster (Tran Ba Huy et al., 1983). Furthermore, sequestration in renal epithelial tubular cells is a causal factor of aminoglycoside nephrotoxicity (Quiros et al., 2011). The persistence and/or sequestration of aminoglycoside antibiotics in the endolymph may augment ototoxicity either through increased structural damage to inner ear hair cells or through mechanisms that ultimately lead to cell death. Therefore, our antibiotic exposure regimen comprises estimated therapeutic treatment concentrations (25 µM) as well as concentrations that have effectively damaged lateral line hair cells in previous work (400 µM: Chiu et al., 2008; Coffin et al., 2013).

Under our experimental conditions, antibiotics delivered at 25-400 µM did not affect larval mortality (100% survival across all experimental groups). During the experimental period (72-96 h), control and treatment group larvae developed from S46/47 to S47/48. Our antibiotic dosage regimen did not elicit discernible differences between the startle responses of control and treatment larvae. The similar developmental progression of animals across experimental groups indicated that the concentrations of antibiotics (25-400 µM) used for these experiments had no detectable effects on larval external morphological development.

SEM images demonstrated concentration-dependent ultrastructural differences between control and treatment neuromasts. It should be noted that we could not discriminate between kinocilia and stereocilia because our SEM analysis does not allow us to determine whether projections comprise a 9 × 2 microtubule arrangement. Therefore, we have referred to “ciliary” bundles which may or may not include a kinocilium throughout this work. Exposure to concentrations of antibiotic greater than 100 µM resulted in a reduction of hair cell bundle number, indicating their potential for ototoxicity (Figs. 1-3). Cilia also appeared shorter in treatment bundles, particularly those from groups treated with higher concentrations of antibiotic (Figs. 1 and 2). However, because the hair cell bundle sterocilia and/or kinocilia appear as coiled strands in SEM images, it was not possible to measure ciliary length. Even though SEM images illustrate that neuromasts have sustained damage to bundle ultrastructure, we observed a normal startle response from treatment group larvae (Figs. 1 and 2). The similar startle response across experimental groups suggests that mechanotransduction was functional in hair cells of the lateral line and/or inner ear even after exposure to antibiotics (25-400 µM) for 72 h. These findings are in agreement with previous research that shows the startle response is partially mediated by inner ear hair cells (Buck et al., 2012; Mirjany et al., 2011; Simmons et al., 2004).

*Xenopus* bioassays that screen industrial chemicals for both teratogenicity and potential to cause endocrine perturbation are well established (Fini et al., 2012; Fort and Bantle 1990; Fort et al., 2011). Our results have expanded the utility of *Xenopus* as a model cellular bioassay for evaluating potential ototoxins. Mechanosensory neuromast structure and function can be compared in a standardized and consistent manner with high resolution imaging protocols such as the one presented here. The advantages of a *Xenopus* model for investigations of chemical toxicity are furthered by the ease with which the genus *Xenopus* can be housed and used to breed thousands of larvae at a time, offering the possibility of high-throughput analyses using the *Xenopus* lateral line. Moreover, humans are closer to *Xenopus* than to bony fish in evolutionary time, and the well-established Woods Hole *Xenopus Resource Center* can facilitate access to genetic stock and enable novel drug development that builds on knowledge of the *Xenopus* genome and the implementation of contemporary approaches such as CRISPR for targeted genetic analysis and manipulation (Horb et al., 2019; Nenni et al., 2019; Pearl et al., 2012). For these reasons, *Xenopus* holds promise as a surrogate species for biomedical investigations of the inner ear and other organ systems in disease (Blum and Ott, 2018; Exner and Willsey, 2021; Ramirez-Gordillo et al, 2015). For example, amphibians such as *Xenopus* afford an opportunity to examine ultrastructural features of hair cell regeneration in the lateral line with imaging protocols such as the one presented here (Groves, 2010).

## 5. CONCLUSION

Our results demonstrate the utility of *Xenopus* as a means for evaluating the impact of water-soluble chemicals such as antibiotics on mechanosensory hair cell ultrastructure. The use of scanning electron microscopy in this study has provided evidence that hair cell bundle damage can be detected at micromolar antibiotic concentrations and adds a high-resolution imaging approach to the screening repertoire that can complement studies of live neuromasts with fluorescence microscopy (Harris et al., 2003; Nishikawa and Sasaki, 1996). Future undertakings with this bioassay at a larger scale could be simplified by using an environmental scanning electron microscope (ESEM), thus reducing the specimen preparation time prior to imaging (Lee and Touissant, 2018; Muscariello et al., 2005). Phenotypic bioassays such as the one presented here can be used to evaluate novel therapeutic compounds for potential side effects, as well as for their otoprotective potential, thereby expediting the drug development process during the discovery phase. Because no one species can serve as proxy for humans, the applicability of research findings from screens using model organisms can be strengthened by comparative studies demonstrating commonalities and differences in the effects of candidate drugs on mechanosensory hair cells from many different species.

## List of Abbreviations

ESEM: environmental scanning electron microscopy
GENT: gentamicin
NEO: neomycin
SEM: scanning electron microscopy
STREP: streptomycin

## Author contributions

EES conceived of the study and supervised the research; VBK, and AL implemented the experimental plan; VBK AL, and EES contributed to the design of the research, to the analysis of the results, and to the preparation of the figures, tables, and manuscript.

## Acknowledgements

We are grateful to Dr. Peter Cooke for technical advice and access to the imaging resources of the NMSU Core University Research Resources Laboratory and thank Rosalinda Corchado and Kimberly Rodriguez for assistance with manuscript preparation.

